# A fast, general synteny detection engine

**DOI:** 10.1101/2021.06.03.446950

**Authors:** Joseph B. Ahrens, Kristen J. Wade, David D. Pollock

**Author notes:** Corresponding authors Joseph B. Ahrens, David D. Pollock.

## Abstract

The increasingly widespread availability of genomic data has created a growing need for fast, sensitive and scalable comparative analysis methods. A key aspect of comparative genomic analysis is the study of synteny, co-localized gene clusters shared among genomes due to descent from common ancestors. Synteny can provide unique insight into the origin, function, and evolution of genome architectures, but methods to identify syntenic patterns in genomic datasets are often inflexible and slow, and use diverse definitions of what counts as likely synteny. Moreover, the reliable identification of putatively syntenic regions (i.e., whether they are truly indicative of homology) with different lengths and signal to noise ratios can be difficult to quantify. Here, we present *Mology*, a fast, flexible, alignment-free, nonparametric method to detect regions of syntenic elements among genomes or other datasets. The core algorithm operates on consecutive, rank-ordered elements, which could be genes, operons, motifs, sequence fragments, or any other orderable element. It is agnostic to the physical distance between distinct elements and also to directionality and order within syntenic regions, although such considerations can be addressed *post hoc*. We describe the underlying statistical theory behind our analysis method, and employ a Monte Carlo approach to estimate the false positive rate and positive predictive values for putative syntenic regions. We also evaluate how varying amounts of noise affect recovery of true syntenic regions among *Saccharomycetaceae* yeast genomes with up to ~100 million years of divergence. We discuss different strategies for recursive application of our method on syntenic regions with sparser signal than considered here, as well as the general applicability of the core algorithm.

## Introduction

When biological objects have shared ancestry, they are said to be homologous. DNA and protein sequence homology can be inferred based on alignment and evaluation of sequence similarity, but over evolutionary time the accumulation of insertions, deletions, and mutations make accurate alignments infeasible. The structural arrangement of sequences along genomes may be retained longer, and can provide independent information about homology and the evolution of genomes. One useful structural evolutionary signal is conserved clustering of genes among divergent genomes, called *synteny*. The term ‘synteny’ was originally used to denote the presence of multiple genetic loci on the same chromosome (Renwick 1971; Nadeau 1989), but now often refers to a common localization or ordering of identifiable genetic elements among genomes that is associated with homology.

Given this relationship between shared ancestry and organization, synteny provides an excellent framework through which to understand genome evolution (Hurst et al. 2004; Oliver and Misteli 2005; Splinter et al. 2006; Batada and Hurst 2007; Michalak 2008; Xu et al. 2009). However, the biological underpinnings driving selection for genome architecture are quite complex and can inhibit the detection of meaningful syntenic signal. Factors that affect the evolution of synteny include effective population size (Lunter et al. 2006; Molina and van Nimwegen 2008; Koonin and Wolf 2010; Weber and Hurst 2011), large paralogous gene families (Fischer et al. 2001; Innan and Kondrashov 2010; Sugino and Innan 2012), genomic scope in terms of macro vs. micro synteny (Putnam et al. 2007; Putnam et al. 2008; Srivastava et al. 2008), presence of structural elements such as nucleosomes and chromatin looping (Oliver and Misteli 2005; Splinter et al. 2006; Batada and Hurst 2007; Spitz and Duboule 2008; Tena et al. 2011; Bagadia et al. 2016), genome rearrangement events (Fischer et al. 2006; Denoeud et al. 2010), intergene distance (Poyatos and Hurst 2007), and variable regulatory constraints acting on adjacent genes (Hurst et al., 2002; Kensche et al., 2008; Wang et al., 2011, Woolfeet al., 2005; Royo et al., 2011; Tena et al., 2011; Irimia et al., 2012).

The biological factors that affect evolution of synteny impact how it should be detected and evaluated. In particular, it is useful to compare synteny detection to sequence alignment, which has the same aim to use similarity to infer homology, but differs in its assumptions of how the underlying elements are structured and how they change over time. In sequence alignment, the number of character states is limited to 4 for nucleotide sequences and 20 for amino acid sequences, while synteny analysis may involve tens of thousands of different genes. Furthermore, the states of nucleotides and amino acids are relatively easy to identify, whereas the gene categories used in synteny analysis may be uncertain. Because of the limited number of states, nucleotide and protein sequences tend to exhibit frequent homoplasy (independent evolution to the same state along different lineages), while recurrent evolution to the same gene category is rare. In contrast, gene order and clustering may be more malleable than nucleotide and amino acid rearrangement within sequences. Because of this, most sequence alignment algorithms assume collinearity (that the ordering of sites in homologous sequences is conserved), while for synteny analysis the collinearity assumption may sometimes be unrealistic (Fischer et al. 2006; Denoeud et al. 2010). Given the markedly different assumptions one can make when comparing large genomic regions, as opposed to sequences within individual genes, there is growing enthusiasm for improved synteny detection methods that scale well for genomic data (Armstrong et al. 2019).

The characterization of synteny is confounded by a multitude of synteny concepts in the literature, many of which are incongruent with one another, pertain to synteny occurring at vastly different scales, and entail different analytical methods. Durand and Sankoff (2003) defined “gene clusters” as sets of genetic loci that are all contained within a short interval on a chromosome. Raghupathy and Durand (2009) then showed that expected frequencies of gene clusters in genomic datasets can be approximated—assuming that the genes are randomly ordered—if the copy numbers of all genes are equal (that is, that each gene occurs exactly 2, 3, 4, etc. times within the dataset). However, calculating the expected number of gene clusters in the general case appears to be computationally intractable (Durand and Sankoff 2003; Raghupathy and Durand 2009). Luc et al. (2002) described an algorithm to identify what they term “gene teams” (or “δ-teams”) which differ from the above gene clusters in that adjacent genes must be within a certain sequence distance (δ), while the entire team may be arbitrarily large. He and Goldwasser (2005) extended this algorithm by removing the constraint that each gene can only occur once on a chromosome, referring to the more general patterns as “homology teams”. Gene teams and homology teams are notably similar to the “max-gap clusters” from pairwise genomic datasets described by Hoberman et al. (2005), who also discuss methods for calculating the expected number of such clusters, assuming a pairwise genomic dataset (where each gene has exactly one matching predicted homolog).

The gene clusters described by Durand and Sankoff pertain to synteny on a small scale (i.e., “microsynteny”). In contrast, the homology teams of He and Goldwassser (2005) may characterize synteny at larger scales, as they may contain any number of genes. However, the relative gene ordering among putatively syntenic homology teams may be completely different, and to our knowledge, there are no easy methods for calculating expected numbers of gene teams, homology teams or max-gap clusters in the general case (Hoberman et al. 2005; Raghupathy and Durand 2009). Many computational tools also exist to visualize putative syntenic regions within genomic data, often including their own particular rules to define synteny (Sinha and Meller 2007; Cai et al. 2011; Cleary and Farmer 2018). Some methods require collinearity, and some do not, but all synteny methods, visual or otherwise, must account for the probability of randomly finding homologous genes on the same chromosome or in the same small region for the genomes compared (Hampson et al. 2003). A popular null model is the assumption that the ordering and matching of homologous genes in a dataset are both random (Raghupathy and Durand 2009).

Here, our aim was to develop a general, fast, and flexible yet rigorous method of synteny detection. We wanted it to function well on large-scale genomic data with millions of elements and hundreds or thousands of species, and to that end we avoided situation-specific parameterizations and time-consuming alignment procedures. We call our implementation *Mology.* Given a set of rank-ordered elements, the *Mology* core algorithm searches for sets of *n* elements clustered within a rank-order distance threshold *d* of each other (a *dn-tuple*) that have a paired *dn-tuple* elsewhere in the dataset. To be a pair, each element in the second *dn-tuple* in the pair must be linked to one of the *n* elements in the first *dn-tuple*. The elements in the second *dn-tuple* must also be within *d* of each other, but not overlapping with the first *dn-tuple*. These links, which we will call matches, are based on predictions, from independent information such as sequence similarity, that a pair or group of elements may be homologous. *Mology* efficiently identifies and reports *dn-tuple pairs* containing these matching elements as possible syntenic regions, while intervening, unmatched elements (noise) are ignored. The choice of *d* allows the scale of a given analysis to range from microsynteny to larger-scale synteny. Consecutive *dn-tuple pairs* from one analysis can, in principle, be merged into a new set of rank-ordered genetic elements, and the core algorithm can be applied recursively to the resulting dataset.

The underlying approach can be applied to ordered elements of any kind, and is thus highly general. All that is required in the case of genomic data is that sequence features (e.g., genes, functional domains, or non-overlapping oligomers) can be reduced to sets of matching (putatively homologous), ordered elements. By operating on rank-transformed element locations, the method is agnostic to the input data type and physical distances among elements. Unique elements, such as single-copy oligomers or novel genes, can be eliminated prior to analysis, in some cases leading to large-scale data reduction and further speed improvement. The statistical framework—and associated null distribution used to estimate probabilities—is nonparametric, making no modeling assumptions about the processes that may have given rise to the putative homologous (i.e., matching) elements, or the downstream events which caused genome architectures to diverge. The null model is based solely on whether the genetic elements appear to be randomly distributed. Below, we describe the core *Mology* algorithm, and a statistical framework in which *Mology* output can be evaluated. To our knowledge, this approach has not been described elsewhere. We simulated datasets to show that there are strong expectations under the null model, and used a noisy yeast protein dataset–where putative homologs were intentionally assigned using overly-sensitive similarity criteria—to illustrate a simple, Monte Carlo strategy for estimating false discovery rates in real datasets with complex distributions of matching elements.

## Results

### The Mology core algorithm

The only required input for a *Mology* analysis is a dataset consisting of rank-ordered elements that can be matched with one another in pairs or small groups. The definition of whether elements match is flexible and largely left to the user, but is meant to link elements that are similar enough that they may be homologous. For example, one might rank-order genes according to their positions in genomes, and match them if their sequences are more similar than some chosen threshold. If there is no natural order in some parts of the dataset (such as among chromosomes, or among scaffolds within a chromosome that have not been placed along that chromosome), those parts can be ordered arbitrarily. Singletons, elements that do not match other elements, may be removed before further analysis, but removal is not required. Because *Mology* works with rank-ordered integers, in principle it can be applied to a wide variety of datasets, including some outside of genetics; however, we will often refer to the ranked elements as *genetic elements* here to maintain our biological example.

As mentioned in the introduction, the core algorithm searches for paired *dn-tuples* in a rank-ordered dataset, with the pairs defined by matches between each of the *n* elements in the first *dn-tuple* with elements in the second, both sets within *d* of each other but not overlapping. For example, if *d* = 10, we might have three elements in a set of rank-ordered gene positions in the range [0,10] paired with three other one-to-one matching elements in a set in the range [1012,1022]; this would be a *d3-tuple pair* with *d* = 10. A d3-tuple pair is a dn-tuple pair with 3 elements in each clustered set, and we will name larger clusters analogously. A visual example of a d3-tuple and d5-tuple are shown in Figure 1. If the number of matching genes, *n*, differs between the dn-tuples in a pair, the pair is characterized by the shorter dn-tuple. In a post-processing stage, overlapping *dn-tuple pairs* are merged into non-overlapping sets, and so we will sometimes refer to the raw *dn-tuple pairs* and paired merged sets as *paired clusters*.

**Figure 1.**
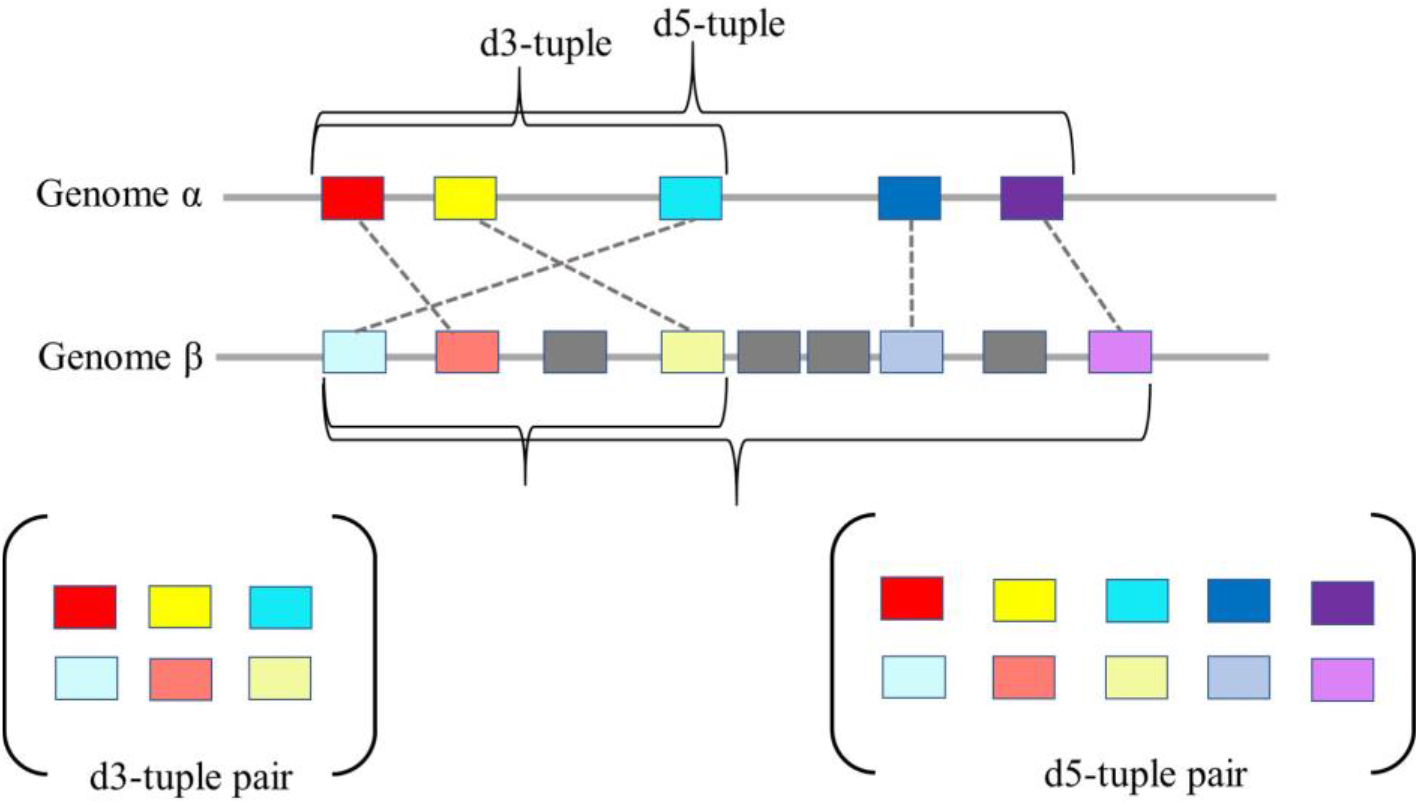
Visualization of a d3- and a d5-tuple pair from matching, ordered genomic data. Colors indicate elements that exhibit enough sequence similarity to indicate possible matches, with the darker shade belonging to Genome α and the corresponding lighter shade indicating a matching element on Genome β. Grey squares indicate Genome β elements that do not match Genome α in this region. If they are singletons in the overall genome comparisons, they can be excluded from *Mology* rank ordering. There are a total of 10 d3-tuple pairs with various unordered combinations of the elements in the d5-tuple pair, and these overlapping dn-tuples would be merged to paired regions of at least length 5 in the post-processing stage.

Raghupathy and Durand (2009) point out that the search space for the *dn-tuple pairs* problem is proportional to *N*^2^, where *N* is the total number of elements in the dataset. A sub-optimal method of finding such pairs is to compare every possible reference window (e.g., [0,10], [1,11]… [*N* − *d*, *N*]) to every non-overlapping window of equal size, remove duplicates, and observe the number of elements shared.

Improving on the above strategy, *Mology* also starts with every possible reference window, but then looks up the complete set of matches to the elements in that window, sorts them, and identifies clusters within width *d* of each other in the sorted set and greater than the reference window elements. In further detail, for a given window with integers {*x*_*i*_… *x*_*i*+*d*_}, if the correspondence set *X*_*j*_ is defined as the integer positions that match the elements in that window, but are themselves outside of that window, then our aim is achieved efficiently by sorting *X*_*j*_. The sorted list is processed linearly to identify all sets of *n* adjacent integers {*x*_*j*_… *x*_*j*+*n*_} such that *x*_*j*+*n*_ − *x*_*j*_ ≤ *d*. Any set identified in this step, along with the matching integer positions in the starting window, forms a dn-tuple pair. The algorithmic complexity thus involves a linear progression of windows along the rank-ordered elements, a non-linear integer sort, and a linear progression through each sorted list. The only non-linear component is the integer sort, which is one of the most highly optimized algorithms/transformations in computer science and very fast. Also, it is not necessary to consider matches with lower rank order than the elements in the starting window because they will have already been identified.

### Simulated data and the null model

The dn-tuple pairs identified in a *Mology* run represent potentially syntenic regions within or between genomes, but to evaluate the evidence for synteny it is necessary to calculate the probability of finding matching clusters by chance alone. We quantify this using the null hypothesis that the elements are ranked independently with respect to their matches. The expected frequency of paired clusters under this null hypothesis depends on the window size *d*, the total number of matched elements *N*, and the copy numbers of all elements (that is, how many elements have 1, 2, 3, etc. matches elsewhere in the dataset). Expectations also depend on the minimum size, *n*, of the paired clusters considered. For a given data set, the window size and cluster sizes considered can be adjusted to control positive predictive values (PPVs, the estimated proportions of clusters that are not accounted for by null expectations, equal to one minus the false discovery rate, or FDR). The analytical needs of different researchers may vary: setting high PPVs (e.g., 99% or above) will help to avoid false positives, whereas setting lower PPVs (50% or less) will detect signals of synteny that may be mixed in with considerable noise. Our implementation of the *Mology* algorithm outputs the PPVs associated with a chosen *n* threshold, and decisions regarding appropriate levels of false discovery are left to the user.

The expected outcomes under the null model can be determined using simulated replicate datasets, with the same distribution of matching elements as the real input dataset, but randomly ordered with respect to matches. We first considered the special case where every element in the dataset has exactly one match (that is, the elements are paired). To implement the simulation, we chose randomly paired integers in the range [0, *N* − 1] without replacement, resulting in *N*⁄2 subsets of integer pairs (Figure 2). When the total number of elements in the dataset *N* is large relative to the distance threshold *d*, the expected number of d2-tuple pairs under the null model is roughly *d*^2^ (see Methods). The expected number of d3-tuple pairs is about *d*^4^⁄*N*, the expected number of d4-tuple pairs is about *d*^6^⁄*N*^2^, and in general for clusters of size *n* and large *N* relative to *d*, the expected number of dn-tuple pairs can be approximated by *Nd*^2(*n*−1)^⁄*N*^(*n*−1)^. This approximation holds well for d2-tuple counts when *d*^2^⁄*N* ≪ 1 (Figure 2; e.g. *N* = 100,000; *d* ≤ 100), which makes sense in that the approximation does not account for overlapping dn-tuples, and overlapping dn-tuples are rare under these conditions. For dn-tuple pairs with larger cluster sizes (*n* > 2), the approximation begins to deviate at smaller *d* values (Figure 2b), but this is compensated by the observation that expectations are quite small and the approximations are always conservative.

**Figure 2.**
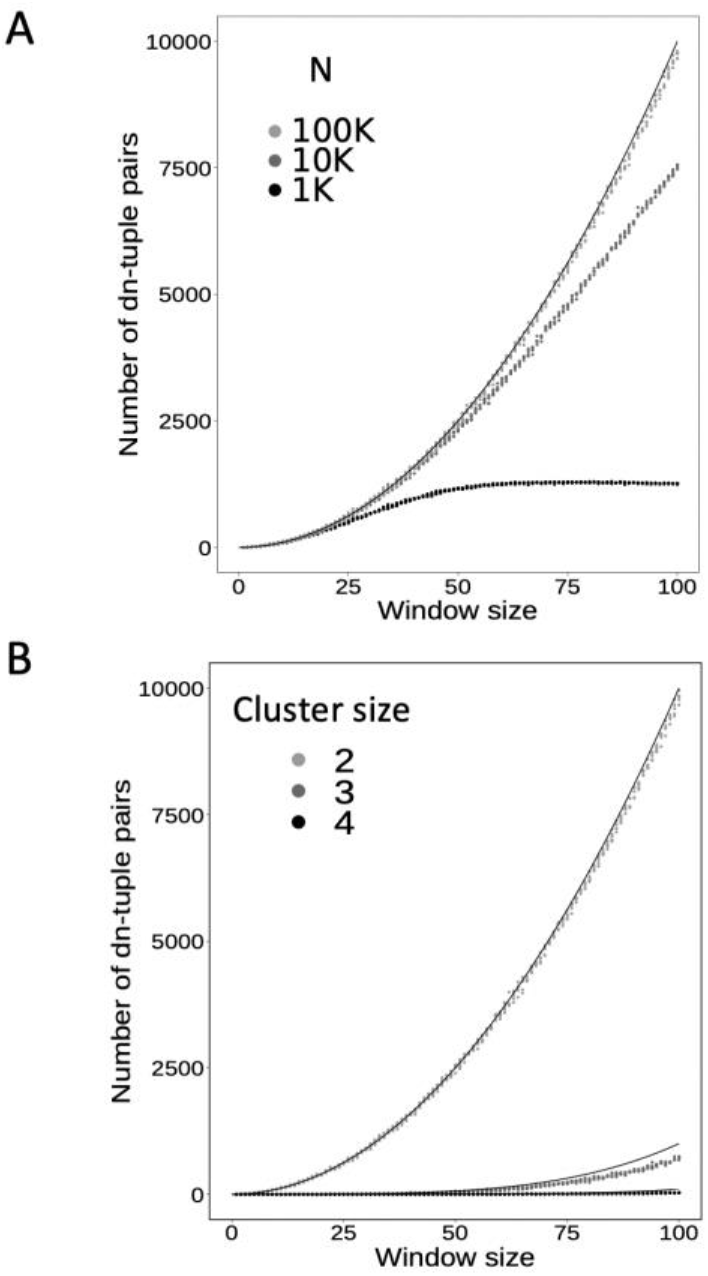
Predicted versus simulated dn-tuple pairs. Dn-tuple pairs were counted for for 5 simulated replicates under each condition, using window sizes (*d*) ranging from 1 to 100 (3,000 simulations overall). A) simulation results for *N*=100K, 10K, and 1K are shown in light, medium and dark grey dots, respectively, run with cluster size *n*=2. B) cluster sizes of *n*=2, *n*=3, and *n*=4 are shown for simulations with a sample size of *N*=100K, and respective prediction functions (*Nd*^2(*n*−1)^⁄*N*^(*n*−1)^) shown as thin solid lines closest to each simulation result.

A practical example of this special case is the set of reciprocal best hits (RBH) in a pair of genomes (Wolf and Koonin 2012; Hernández-Salmerón and Moreno-Hagelsieb 2020). Although this special case is analytically tractable, the general case with complex distributions of matching elements is not considered tractable (Durand and Sankoff 2003; Raghupathy and Durand 2009) so it is useful to compare the simulation results to analytical expectations for this case. For most datasets with more complex and realistic distributions of matching elements, we must rely on fast simulations to obtain more accurate PPV estimates. However, this special case helps to illustrate the trade-offs in considering workable values of *d* and *n* for a given dataset.

### Monte Carlo estimation of false discovery rates across seven yeast genomes with noisy signals

A limitation of many current synteny algorithms is their susceptibility to highly divergent genomes, and also to incomplete, poorly annotated, or noisy datasets. To evaluate *Mology’s* performance in the presence of noise, we used simplistic similarity-based homolog annotations that identify some but not all matching protein-coding sequences in genome datasets, and that sometimes produce likely false matches based on random similarity. Our fast, intentionally noisy protein homolog prediction algorithm is based on kmer (substrings of length *k*) matching to protein sequences of a reference species. *Saccharomyces cerevisiae* was chosen as the reference since its extensive characterization make it an ideal model for method validation. This similarity detection algorithm identifies proteins with multiple matching in-frame amino acid kmers of a given length within a given sequence distance *w*. We did not optimize for sequence homology detection but were able to easily control the noisiness or stringency level by altering the length of kmers used and other algorithm parameters (Table 1).

**Table 1.** Protein-coding similarity parameters to produce varying levels of background noise, from most noise (Stringency 3) to least noise (Stringency 1).

| Parameters | Stringency 3 | Stringency 2 | Stringency 1 |
| --- | --- | --- | --- |
| k | 4 | 5 | 5 |
| Non-overlapping window size (bp) | 25 | 50 | 25 |
| Minimum kmer matches per window | 4 | 5 | 5 |
| Maximum number of windows separating adjacent matches | 2 | 3 | 3 |
| Minimum number of adjacent windows with hits | 2 | 2 | 2 |
| Do genomic kmer matches have the same order as the reference protein? | Yes | No | Yes |

We applied this similarity mapping algorithm to sets of potentially homologous protein fragments (i.e. matches) among seven well-annotated *Saccharomycetaceae* yeast species: *Saccharomyces cerevisiae, S. paradoxus, S. kudriavzevii, Kazachstania africana, Tetrapisispora phaffii, Zygosaccharomyces rouxii* and *Kluyveromyces lactis* (we will call this dataset Y7G). This yeast family underwent a whole genome duplication (WGD) ~75 MYA, meaning that there is both inter-species and intra-species synteny. The first step in this analysis was, for each of the seven species and each stringency level, to generate the set of all genes with high similarity to *S. cerevisiae* protein-coding sequences (Supplementary Data 1). These are organized into matching groups consisting of the subset of sequences that matches each *S. cerevisiae* protein-coding sequence (Supplementary Data 2). This was done separately for each stringency. The total identified reference (*S. cerevisiae)* coding sequences for each comparative species, at each stringency, can be found in Supplementary Table 1.

To assess how much of the yeast genomic *Mology* output is meaningful signal, pseudoreplicates of the Y7G dataset were generated by randomizing the gene integer positions in the original file (see Methods). For the Y7G dataset and each pseudoreplicate, we used *Mology* to detect dn-tuples at *d* = 10, for values of *n* from 2-10 (Table 2, Supplementary Data 3, Supplementary Data 4). For pseudoreplicates, the average counts are shown for 100 replicates independently generated for each set of parameter combinations. For larger numbers of clustered elements *n*, the number of dn-tuple pairs expected by chance under the null model becomes vanishingly small compared to the number of pairs in Y7G, with clear conservative cutoffs of about one or fewer pairs expected by chance for runs when the number of matching elements in the dn-tuple pairs were *n*=4, *n*=5, and *n*=8 for stringency 1, 2 and 3, respectively.

**Table 2.**
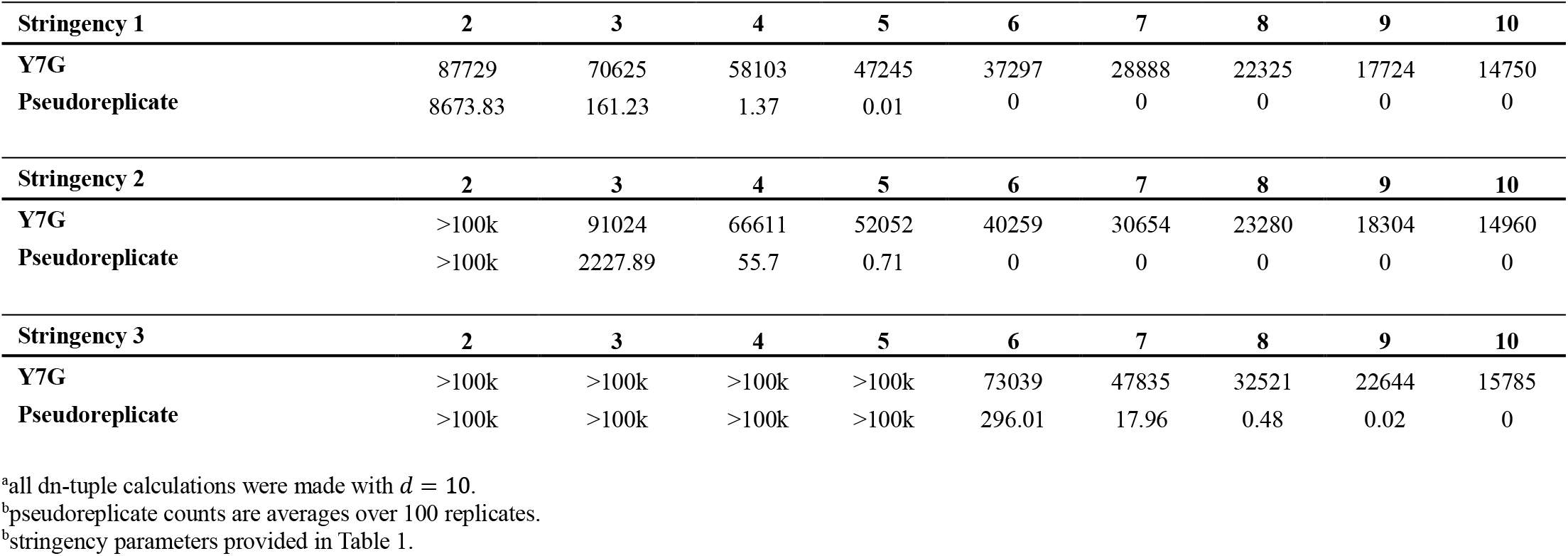
Counts of dn-tuple^a^ pairs in Y7G and corresponding pseudoreplicates ^b^, at three stringency levels ^c^.

We note that the number of dn-tuple pairs detected in these analyses, in both real data and simulations, were for dn-tuples found in all window comparisons; these dn-tuple pairs likely have overlapping element membership. This is not a problem here above the conservative cutoffs because the probability of chance overlaps is small and accounted for in the pseudoreplicates. However, it is more straightforward to interpret results for independent paired regions, so in subsequent analyses we calculated counts, expectations, and FDRs for continuous merged regions, or CMRs (Figure 3; Supplementary Table 2). The CMRs are simply the merge of all overlapping paired dn-tuples and their length was characterized based on the number of matched elements in a given merged region of a pair (Supplementary Table 2, Supplementary Data 5, Supplementary Data 6). Because many CMRs are much longer than expected, we also calculated corrected expectations and corrected FDRs for a given CMR length based on removal of PPV probable elements from *N* in the null model calculated from longer CMR pairs (see Methods). These refined estimates are calculated under the hypothesis that elements that are probably syntenic should be removed from the null model calculations.

**Figure 3.**
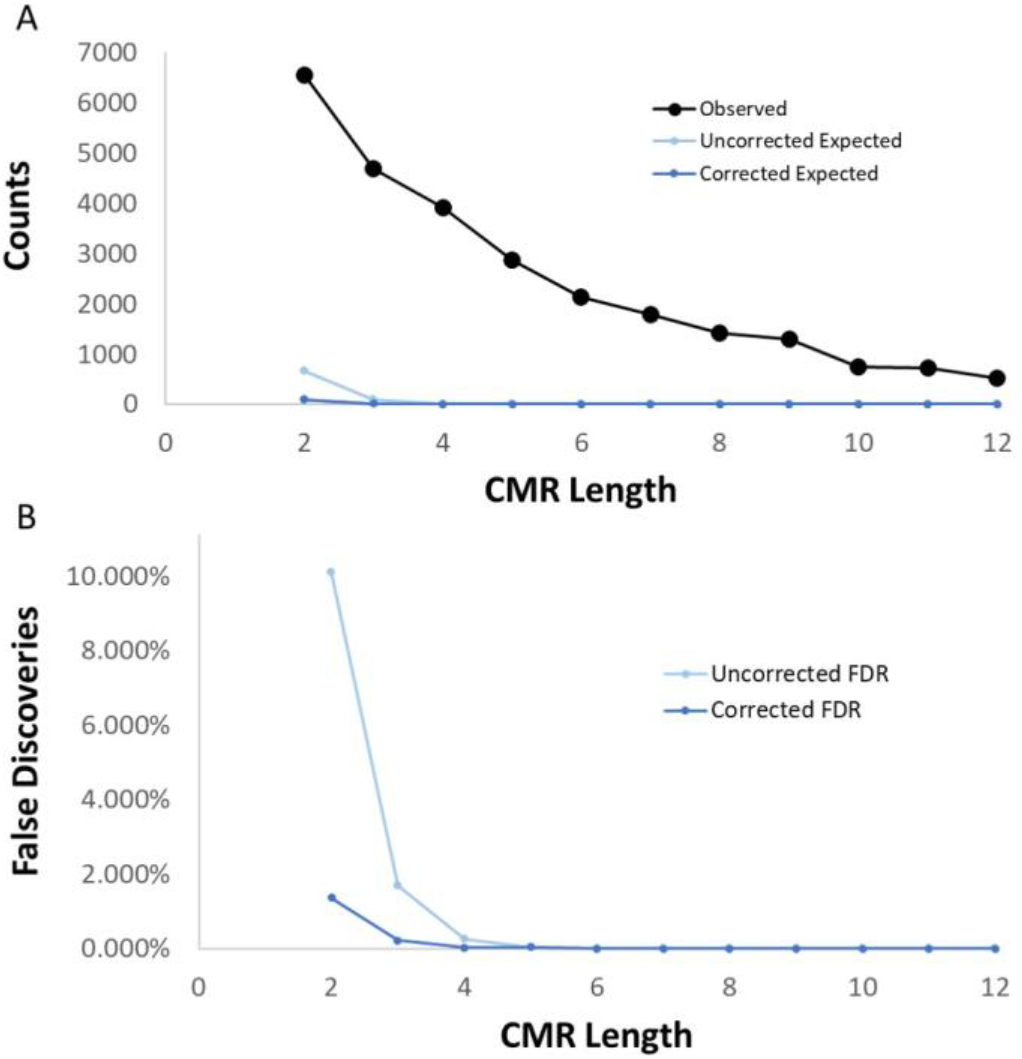
*S. cerevisiae* self-self comparison CMR element counts and predictions. CMRs were identified based on overlapping *Mology* results applied to Y7G with stringency 1 match predictions with *d=10* and *n=2*. Counts are shown for the total number of elements in CMRs greater than or equal to the given length, and uncorrected expectations were obtained by accounting for the self-self comparison as a fraction of all genome comparisons. Corrected expectations were obtained by excluding the proportion of elements that were already identified as a homologous match (see Methods). A) Observed, uncorrected expected, and corrected expected are shown in black, light blue, and blue, respectively. B) Uncorrected and corrected FDRs are shown in light and dark blue, respectively.

We ran *Mology* on the Y7G dataset with *d* = 10 and *n* = 2 to *n* = 10 (Supplementary Table 2), but to illustrate the CMR results it seemed most relevant to focus on the *n* = 2 dataset at stringency 1, as this setting has the most confident (least noisy) coding sequence predictions and encompasses the regions identified using higher values of *n*. We also focused on the *S. cerevisiae* genome self-self (or intra-genome) pairings because the bulk of matching regions in this comparison should have arisen from the ancestral whole genome duplication (discussed in further detail below). The expected element counts are only a small fraction of the observed element counts for all CMR lengths (Figure 3a), and the corrected FDR is less than 2%, even for CMRs of length 2. Whether corrected or uncorrected, essentially all the CMRs of length 4 or more (and maybe 3 or more) are expected to be true positives, and the expectation of spurious predictions is negligible (Supplementary Table 3). The number of likely positively predicted elements in CMRs of length at least two or at least three is comparable to the number of genes in the *S. cerevisiae* genome, which is compatible with identification of synteny for most of the whole genome duplication, with perhaps some further duplication of chromosome segments. The difference between genomic CMRs and the null expectation is similarly distinct for the pairwise comparison between the six other genomes and *S. cerevisiae* (Supplementary Table 2).

One caveat in distinguishing homology from random noise is that under some conditions the post-run analysis can be slowed by the large number of random short dn-tuple matches, so it can be useful to employ a filter prior to merging to avoid the pairs least likely to be actual homologs. It is also worth noting that the true proportion of elements matching due to chance is likely low in the most stringent analysis, and we expect that the true proportion is closer to the corrected than uncorrected FDR values.

However, for less stringent analyses where we create conditions that are intended to include noise, the true proportion will be somewhere in between the corrected and uncorrected FDR values. Regardless, we infer that with appropriately controlled parameter choices *Mology* can be used to reliably identify meaningful syntenic regions, even among smaller gene clusters.

To evaluate the impact of varying the search window parameter, *d*, we also generated dn-tuples and CMRs from a smaller dataset containing only three highly diverged yeast genomes, *S. cerevisiae, Z. rouxii, and K. lactis* (Y3G). This was performed at values *d* = 5, *d* = 10, and *d* = 15 (Supplementary Table 4, Supplementary Data 7). In comparing results between Y3G and Y7G (Table 2 and Supplementary Tables 2-3), the null expectations becomes negligible at similar values of *n*, suggesting that inferences are relatively robust to dataset size. For the Y3G CMRs at stringency 1, null predictions are consistent across the three different *d* values, dropping off below 1% around *n* = 3 or across the three window sizes. For stringency 2, null predictions drop off only slightly later, requiring *n* = 3 to drop below 1% for *d* = 15. For stringency 3 at *d* = 15, *n* = 9 is required but only *n* = 5 at *d* = 5. Notably, the number of genes detected around these cutoffs are not greatly different. These results are not surprising, in that with greater noisiness there will be more random matches in between the truly homologous matches, and window sizes need to be longer to find the same run of true matches.

To illustrate how synteny predictions unfold in a genomic example, we focused on describing a duplicated region that was split between two *S. cerevisiae* chromosomes, VIII and XVI, and contains the TOM70 and TOM71paralogs. We identified this locus in *Mology-*predicted syntenic regions within *S. cerevisiae* self-self comparisons, as well as regions in *Z. rouxii* and *K. lactis* that matched the *S. cerevisiae* copies (Supplementary Table 5).

We compared the *Mology*-constructed region between *S. cerevisiae* and two species that diverged before the whole genome duplication (Figure 4 and Supplementary Table 6). Relative positions in each genome were validated by comparison to the manually curated yeast synteny database, YeastGenomeOrderBrowser (Byrne and Wolfe 2005). At all three levels of annotation noise, dn-tuple pairs were matched to each other correctly, i.e. pairings were made between homologs in the correct order and at accurate regions of the genome (Figure 4). Even against a high level of background noise and intervening false homolog annotations (Supplementary Table 6), a majority of the syntenic regions predicted with the high stringency analysis were still recovered in the noisy data. The inclusion of noisier annotation data enhanced the *Mology*-predicted synteny map by incorporating a few more genes than were matched by the most stringent analysis (Figure 4). Although some likely homologs were not matched, even in the least stringent analysis, the syntenic regions were robustly predicted. A boost in sensitivity is especially important when comparing highly divergent genomes, where sequence similarity may have significantly degraded over time (a common problem in comparative genomics). A sensitive method is also important when mapping duplicated syntenic blocks to pre-duplication blocks because of the complementary loss of genes (see also Figure 4). Overall, these results indicate that *Mology* is well equipped for use in such scenarios, and that it could be used in analyses that allow moderate uncertainty in homology predictions in the sequence comparison stage.

**Figure 4.**
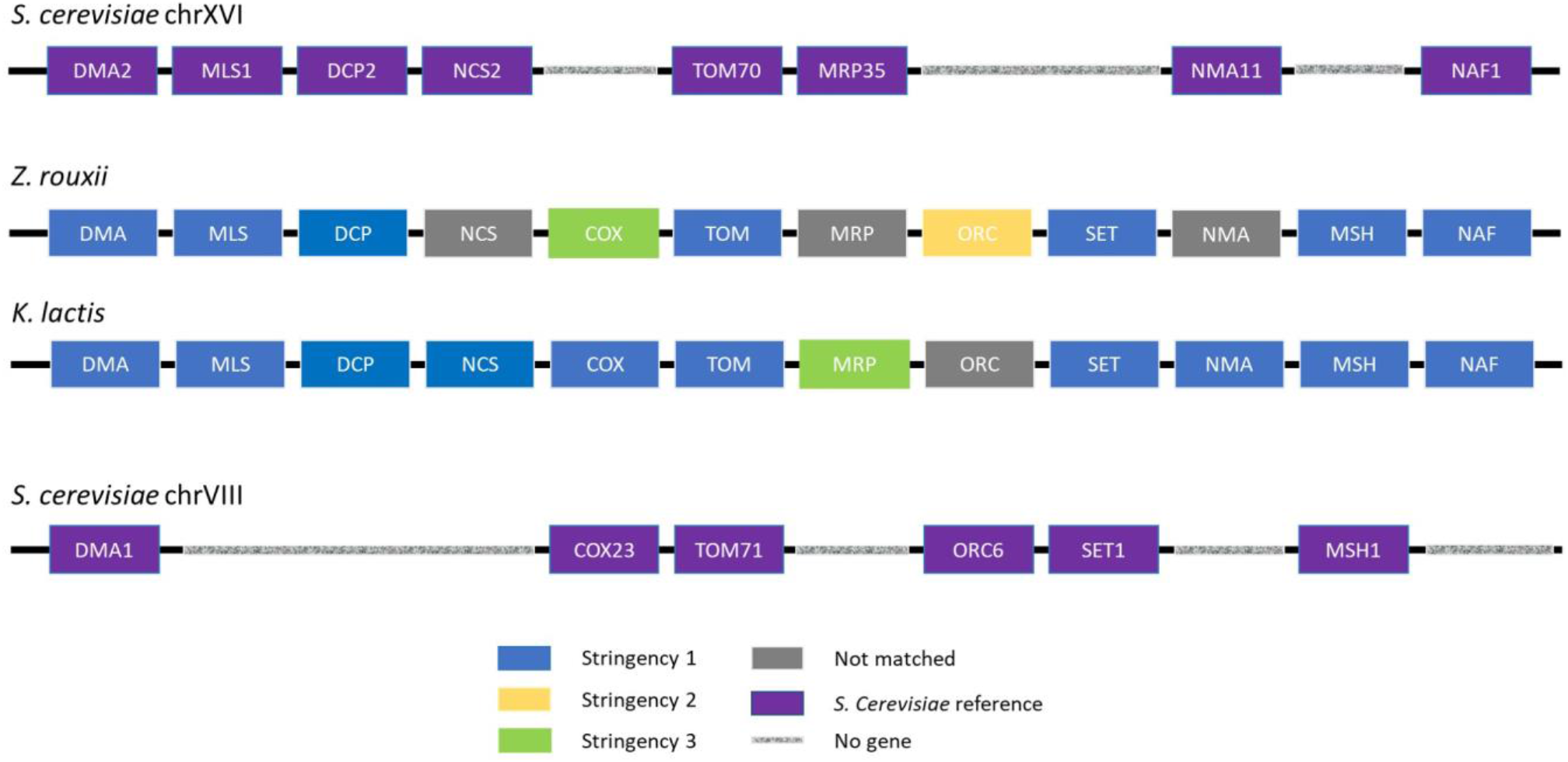
*Mology* results for known syntenic architecture in yeast genome regions containing duplicated Tom70/TOM71 paralogs. Ancestral, pre-duplication chromosomes of *Z. rouxii* and *K. lactis* were identified in syntenic CMRs, with respective matches to the corresponding, post-duplication chromosomes in *S. cerevisiae.* Blue, yellow and green boxes indicate the least sensitive stringency at which homologs were identified. Grey boxes indicate protein-coding sequences that were not matched under any setting using the kmer sequence similarity algorithm. Purple boxes are annotated in the *S. cerevisiae* reference genome, and hatched grey lines are positions where there is no annotated gene.

To further evaluate how synteny decay manifests at the genomic-scale in divergent species comparisons, we evaluated distributions of coding sequences in CMRs paired between all combinations of pairwise Y7G species comparisons at the most stringent annotation setting (Supplementary Table 7). We focused only on pairwise comparisons between *S. cerevisiae* and each other species, as these numbers directly represent shared putative homologs to the reference protein coding sequences. We considered coding sequences in a given genome that were contained within at least one CMR that had a paired CMR within the *S. cerevisiae* genome. For comparison, we also show the total number of genes annotated by the Hittinger group (Shen et al. 2016; Opulente et al. 2018), with 5912-6339 genes in the four post-duplication species and 5338-5430 genes in the two post-duplication species, a mean 13% increase following duplication; and b) the total number of *S. cerevisiae-*matching sequences in each genome (predicted homologs from the sequence comparisons prior to *Mology* analysis). The total number of matches fell from 5454 genes in the closest genome, *S. paradoxus*, to around 3117 in the most distant genome, *K. lactis*. For the *n* = 2 *Mology* analysis, the number of genes in predicted syntenic clusters was consistently close to the mean 86% of the total matches for all genome comparisons (83-89%, with no clear phylogenetic signal or difference across the WGD event; Figure 5). For the two species in genus *Saccharomyces* with *S. cerevisiae* (*S. paradoxus* and *S. kudriavzevii*), there is almost no drop in the number of genes in CMRs going from *n* = 2 to *n* = 7, indicating that syntenic content prediction is insensitive to this parameter for low levels of divergence. For the other four more divergent species, there is a greater drop in the number of genes in CMRs compared to *n* = 2 with each increase in *n* going from *n* = 3 to *n* = 7 (Figure 5 and Supplementary Figure 1), indicating that fewer syntenic genes are correctly predicted with higher *n* values. This is not surprising, although it is perhaps surprising that the drop is not sensitive to phylogenetic distance across this range of four species, and notable that the drops are greatest for the post-WGD species *T. phaffii* (Figure 5 and Supplementary Figure 1). This is perhaps an interesting point for future analysis, but the key point is that the drop-off from *n* = 2 to *n* = 3 is marginal for all species comparisons. Recalling from Figure 3, this is consistent with the corrected FDR estimate of less than 2% for the *n* = 2 high-stringency *Mology* analysis across the entire Y7G comparison. We also leave for future analysis consideration of the question whether more accurate sequence-based synteny detection approaches are worth the time they may cost, and whether a trade-off might be found with speed and accuracy at different stringency levels for whatever pre-*Mology* matching algorithm is used. These questions will only be reasonably answered through detailed analysis of many genomes from a broad range of biological diversity, structural rearrangement, and sequence divergence. Nevertheless, the speed and conceptual flexibility of *Mology* should make such specialist analyses possible. Together, these findings align well with current knowledge of synteny levels among these species (Feng et al. 2017), and strongly support the capacity of the *Mology* analyses to correctly identify synteny.

**Figure 5.**
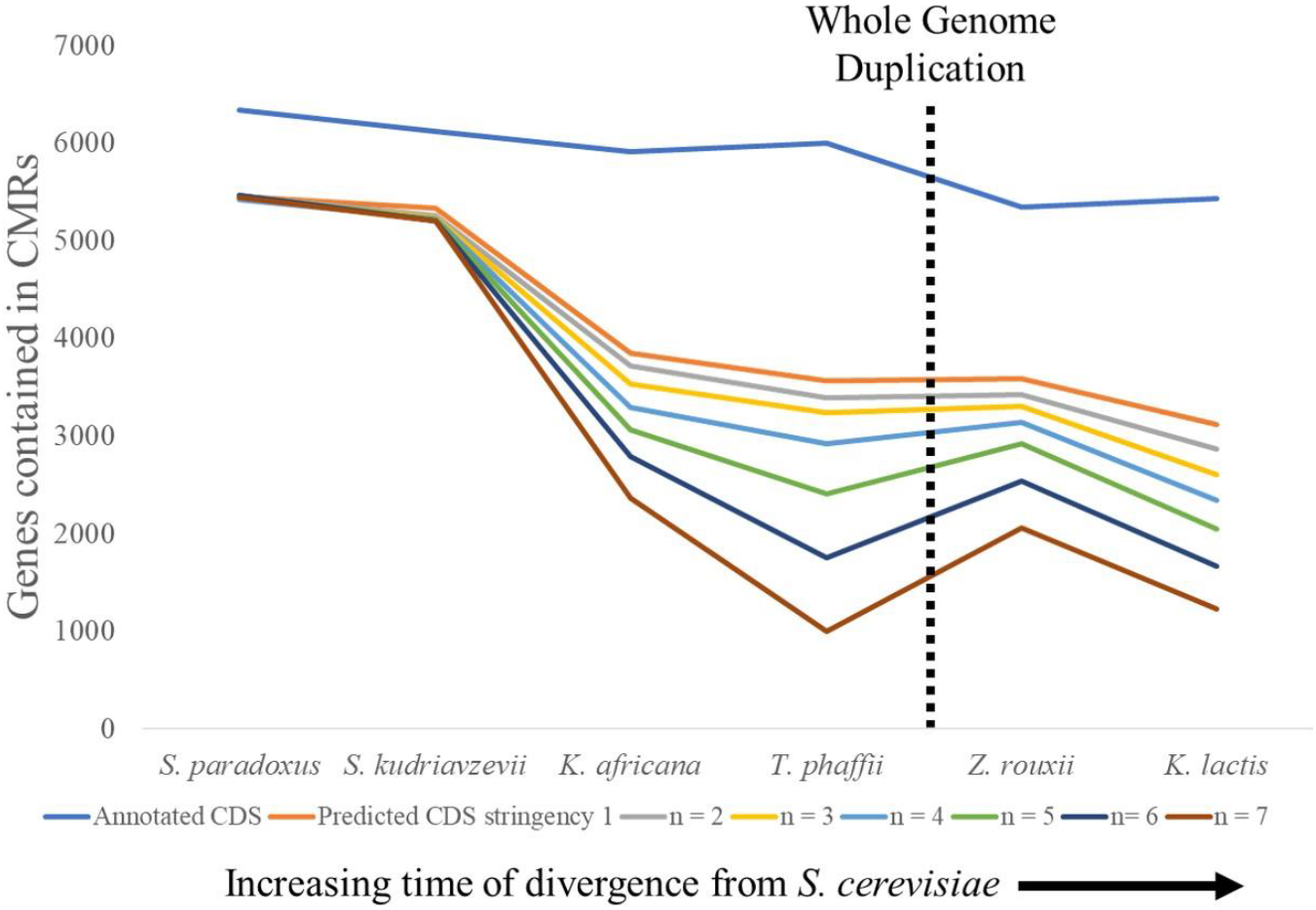
Decay of genes in predicted syntenic regions with *S. cerevisiae* across *Saccharomycetaceae*. The number of genes shown are contained in CMRs between *S. cerevisiae* and six other species from across the budding yeast sub-phylum, Saccharomycetaceae. The number of genes (or CDS) annotated in each genome by the Hittinger group (Shen et al. 2016; Opulente et al. 2018) and by kmer matching are also shown for reference Supplementary Table 6). The CMRs were merged from dn-tuples at each value of n, from 2-7, as indicated in the legend. Species names for each genome are organized from left to right with increasing time of divergence from *S. cerevisiae*. The approximate time of whole genome duplication on this scale is indicated with a dashed black line.

## Discussion

We chose here to use the most direct means to determine the null expectations for paired gene clusters, using randomized pseudoreplicate datasets. Because *Mology* is so fast (Supplementary Figure 2), an analytic solution to the statistical significance problem is, practically speaking, not necessary. For moderate-sized ****N**** (****N**** ~ 100,000) the null distribution can be estimated from ~1000 pseudoreplicates in a matter of seconds per replicate. Raghupathy and Durand (2009) noted that the mathematical framework they developed becomes computationally intractable when dealing with datasets of arbitrary structure (i.e., different copy numbers among different groups matching elements), and an efficient algorithm for identifying gene clusters is, to our knowledge, not described elsewhere. It is also worth noting that ****N**** refers to the number of integers with one or more matches, and large datasets with many unmatched elements can be greatly reduced by eliminating singletons prior to the *Mology* run, enabling analyses that are still quite rapid.

The detection of syntenic regions in molecular datasets is confounded by numerous conceptual difficulties. Currently, there is not a broad consensus regarding i) the formal definition of “matching” elements (Liu et al. 2018), ii) a practical distinction between “synteny” and “collinearity” (Tang et al. 2008; Wang et al. 2012) or, by extension, iii) the significance of syntenic blocks identified in molecular data (Irimia et al. 2012; Liu et al. 2018). Considerable efforts have been made to generalize older, more restrictive notions of synteny (He and Goldwasser 2005; Jahn 2011). Mology takes an approach that is quite flexible, and is capable of rapidly finding unorderered clusters at a microsyntenic level (e.g., spacing between elements of 10 or less), as well as incorporating collinearity in sequences of microsyntenic clusters.

The nonparametric nature of the *Mology* input file can facilitate analysis of a wide range of molecular datasets. The input elements do not need to be genes and element matches do not need to be made with certainty. The element definitions and criteria for matching are up to the end user, provided that the input data can be reduced to sets of well-ordered, matching elements. A fundamentally identical analysis framework can be used, for example, to locate segmental duplications within a single genome, or clustered functional domains within viral polyproteins.

As an example of the conceptual flexibility of *Mology*, one can consider the potential to identify syntenic regions in recursive fashion, allowing access to different levels from micro- to macro-synteny. Consider that the significant (high PPV/low FDR) matching regions identified in one’s initial analysis may be transformed into a new input file by treating non-overlapping CMR pairs as new matched elements. Another strategy might be to remove identified CMRs from an analysis, and then re-run the *Mology* program on the remaining elements. This could identify ancient synteny that had been disrupted by the insertion of chromosomal fragments after divergence. Thus, the core algorithm may perform recursive synteny analyses, allowing the user to identify similarities among genomic architectures at arbitrarily high levels.

The uncorrected method of FDR/PPV estimation implemented in *Mology—*the generation of randomized pseudoreplicate datasets—constitutes a null model in which all elements in the dataset are independently assorted. It should be noted that real datasets may contain long syntenic regions, meaning that by definition a large fraction of the matching positions in the dataset are not independently assorted, so the space of possible random matches is smaller than a Monte Carlo simulation would suggest. We applied a corrected FDR estimate that removes CMR elements from consideration in the null model estimates according to their PPVs, and found that it reduced the FDR for stringent short length (n=2) CMRs by over 5 fold, to below 2%. An alternative probability estimation strategy for short CMRs is a recursive analysis, as suggested above, removing the lowest-probability CMRs and performing a new analysis on the remaining elements from the original dataset. This should have a similar effect in providing more realistic estimates of FDR/PPV for syntenic regions among the remaining positions.

Another key point to emphasize is that the *Mology* core algorithm identifies order-independent dn-tuple pairs. That is, the elements in a dn-tuple do not necessarily occur in the same order in a matching dn-tuple. Users may wish to explore the benefits of considering large *d* values, requiring longer CMRs (subsequent CMR creation is not part of the core algorithm), and interpreting modified probability calculations based on the degree of ordering within the CMRs. Given an estimated FDR in a particular analysis, a perfectly ordered dn-tuple pair would occur by chance with probability *FDR*⁄*n*!. The probability of similar but not perfectly matching orders between pairs can be calculated with more complex formulae. Hence, long ordered dn-tuple pairs and CMR pairs likely represent highly significant syntenic regions beyond the initial *Mology* probability calculation.

The null model used in *Mology* may be inappropriate if elements in the original dataset partially overlap. For example, matching kmers, such as in our *ad hoc* protein similarity identification algorithm are not sequentially independent if they overlap. They will thus form spurious dn-tuple pairs more often than expected under a null model that assumes independent assortment. Methods have been proposed (Ma et al. 2002) to diminish the statistical dependence of matching kmers, such as restricting the input dataset to non-overlapping kmers. Another possibility in this situation is to *post hoc* merge overlapping kmers into single elements or a series of non-overlapping elements if they are long enough, and modifying the lengths and thus the probability calculations of CMRs to adjust to this modification. The *Mology* approach identifies any deviations from the null model, and users should ensure that putatively significant deviations do not have trivial causes, filtering them in the pre- or post-analysis stages if they do.

The *Mology* source code is designed to interact with other programs on the front and back ends, so that users familiar with the *Go* programming language can easily extend functionality as they see fit. They can do this by adding functions to the package file and incorporating appropriate flags in the main program file. Potential applications of *Mology* that transcend genome synteny analyses include an alignment-free statistical framework to identify potential homologous sequences in databases, repeat sequence analysis using clusters or clouds of related repeat-derived kmers (Gu et al. 2008; de Koning et al. 2011; Shortt et al. 2020), and even article keyword matching and other forms of natural language processing. Nonetheless, with synteny being our prime example, we demonstrate here that our implementation of *Mology* constitutes a fast, nonparametric, scalable method for data reduction and general pattern detection.

## Methods

### Input preparation

Given a dataset that appears applicable to *Mology*, pre-processing is required to order the elements and match them into small groups. For the genomic analysis presented here, the gene elements used were CDS annotations from the Hittinger group (Shen et al. 2016; Opulente et al. 2018), and were ordered based on their positions in the *Saccharomycetaceae* yeast genomes that contained them. For joint analyses, chromosomes and the genomes themselves were ordered arbitrarily by chromosome number and by distance from *S. cerevisiae*. The genes in this gene set did not overlap, so we did not need to correct for that, and we made no adjustments for the lengths of genes or the separation of genes along the genome. Singletons, or genes that did not match other genes, were removed prior to further analyses, but they only occurred in *S. cerevisiae*, as all other genomes inherently required a matching sequence in *S. cerevisiae* to be predicted.

In addition to ordering, the genetic elements must be matched into small groups. For the yeast analyses, we applied a low-accuracy kmer-matching approach that is described below. The controllable inaccuracy of the approach allows us to demonstrate that the matching does not need to be correct or certain, and in many cases considerable noise may be tolerated. Once matching and ordering are determined, they need to be converted to the required input for *Mology*, a text file with integer values on each line representing groups of matching elements, and the integer values representing the order of the elements. If singletons are removed, each line will consist of two or more integers separated by space characters. We note that it is necessary for the user to retain a map from the input integers back to any identifiers such as gene IDs that will be needed for post analysis interpretation of *Mology* results. We note that all references to Supplementary Data refer to our file with descriptions of supplementary tables and large raw data files that are too big for the 10 MB supplementary table and figure limits, and which are generally summarized in the main and supplementary tables and figures hosted by the journal; the numbers that follow are helpful guides to where the information is contained in the Supplementary Data file.

### Algorithm and implementation

The core algorithm presented here, which we call *Mology*, was implemented in the *Go* programming language (a.k.a. *Golang*; The Go Authors, 2016). The source code is organized into 1) a *molutils* package containing required functions and data structures, and 2) a *Go* source file, *gomology*, that accepts user input (files and parameter settings) and calls code in *molutils* to produce output. We also include a collection of accessory scripts used in the results presented here to extend and interpret the core output. The codebase is freely available on GitHub (https://github.com/PollockLaboratory/Mology) under the MIT license (https://opensource.org/licenses/MIT).

Initially, *Mology* takes the input dataset *D* containing *N* different integers associated into matching groups, and rank-transforms the *N* integer positions into a modified dataset *D*′. It then creates a map *G*′ relating each rank-ordered position in *D*′ to the rank-ordered position in *D*′ of the matched elements that it was associated with based on co-occurrence in the lines of the input text file. We note that it is a design feature of *Mology* that a rank-ordered integer position may be found in more than one group, and often were, in our yeast analyses, because of the inaccurate protein homology ascertainment method used.

*Mology* proceeds by mapping each set of ranked integer elements in a window of width *d* to every other non-overlapping window that contains elements that match the elements in the starting window. For a given window with integers {*x*_*i*_ … *x*_*i*+*d*_}, if we define the correspondence set *X*_*j*_as the integer positions of all non-overlapping elements that match the elements in that window, then our aim is achieved efficiently by sorting *X*_*j*_. The sorted list is processed linearly to identify all sets of integers {*x*_*j*_ … *x*_*n*_} such that *n* − *j* ≤ *d*. Any set identified in this step, along with the matching integer positions in the starting window, forms a dn-tuple pair. The algorithmic complexity thus involves a linear progression of windows along the rank-ordered elements, a non-linear integer sort, and a linear progression through each sorted list. The only non-linear component is the integer sort, which is one of the most highly optimized algorithms/transformations in computer science and very fast.

Given *D*′ and *G*′, the algorithm proceeds as follows:

1. Define a window [*w*, *w* + *d*], starting with *w* = 0 and *d* a distance threshold specified by the user.
2. Using *G*′, look up all matches *M*_*w*_ to the set of rank-ordered integers *W*_*w*_ that are found in the window [*w*, *w* + *d*], and for which the matches have integer values greater than *w* + *d*.
3. Sort *M*_*w*_ into an ordered set, 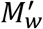.
4. Using the cluster size *n* specified by the user, move in order through 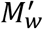 to identify any clustered matches of size *n* that are within *d* of each other. Such matches are by definition dn-tuples that have a paired dn-tuple in the window [*w*, *w* + *d*].
5. Record each unique dn-tuple discovered in a record ({*x*_1_ … *x*_*n*_}, {*y*_1_ … *y*_*n*_}), where *x* and *y* are the ordered elements of the dn-tuple pair that are contained in *W*_*w*_ and 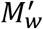, respectively.
6. Increase *w* by 1 and repeat step (1) – (5) recursively until there would be insufficient elements in 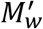 to form a dn-tuple. That is, cease when *w* + *d* > *N* −.

We note that this algorithm could in principle be improved by using windows that overlap by less than *d* − 1, and by simultaneously collecting all dn-tuples for all *n* ≤ *d*. However, the algorithm considered is fast and simple. The most complicated calculation involved is the extremely fast integer sort; the size of each sort involved will depend on *d* and the distribution of the number of matches greater than *w* + *d*.

What we call the core *Mology* algorithm is complete with the output of the complete set of dn-tuple pairs for a given *n* and *d*, sorted by the lowest rank-ordered integer in the pair. In addition, we included Python scripts to merge overlapping paired dn-tuples into stretches we call continuous merged regions (CMRs). The CMRs are categorized by their length.

### Expectations from the null model

Because *Mology* is applied to rank-ordered integers grouped into matched sets, the appropriate null model is that matched sets are created randomly. In this null, the genetic elements are considered to be statistically independent, meaning that deviations from the null model can potentially occur for any non-independent quality, including order homology but also including for example overlap of genetic elements, should it exist. For the example in which all integers are paired into mutually exclusive groups of two, null sets were created by randomly pairing collections of 1000, 10000 or 100000 integers. The yeast datasets were more complex, with overlapping groups of various sizes, and for these datasets the null model was simulated by randomizing the placement of the matched gene integer positions in the original file. In other words, in the starting file for each simulation replicate, each of the *N* integers were replaced with a random integer from 1 to *N*, without replacement.

To calculate analytical expectations or their approximations under the null model where it is possible, we consider that a collection of well-ordered genetic elements with a limited number of matches among themselves—the input dataset required by *Mology*—can be formally conceptualized as a vertex-labeled graph (collection of nodes/vertices connected by edges) with *N* vertices—where the vertices are labelled with all integer positions in the range [0, *N* − 1] (if the position indices begin at 0) and the edges denote matching vertices (where matching vertices may be e.g. known homologous genes or similar sequences). Provided that the individual genetic elements can be considered statistically independent—which should generally be applicable to non-overlapping genetic elements—a reasonable null model is that all elements in the dataset are randomly matched. That is, for any element *i*, the probability that any other element *j* is a match to *i* is exactly equal to *k*_*i*_⁄*N*, where *k*_*i*_ is the number of elements in the dataset that match *i* (i.e., the number of edges connected to *i*, also called the degree of *i*). In the language of graph theory, this null model implies that one may know ahead of time the total number of vertices in the graph *N* as well as their degree distribution (the set of values of *k*_*i*_ for all *i* ∈ [0, *N* − 1]), but that the graph itself is random.

To better illustrate the properties of datasets generated under the null model, let us consider a simple example dataset, in which each element matches exactly one other element. This scenario, where *N* is implicitly even and there are exactly *N*⁄2 edges, is known in graph theory as a perfect matching, and for any set of *N* ≥ 2 labeled (e.g., ordered) vertices, there are (*N* − 1)!! possible perfect matchings, i.e. the semifactorial (*N* − 1) ∗ (*N* − 3)…∗ 3 ∗ 1. Thus, the degree distribution of a random perfect matching on *N* vertices is trivially known (*k*_*i*_ = 1 for all vertices) but other properties of interest follow probability distributions. For example, the expected number of dn-tuple pairs—in a perfect matching of *N* vertices****—****follows a probability distribution determined by the pair of values {*d*, *n*} where *d* is the maximum integer distance between the largest and smallest position in each collection of *n* elements in the pair.

Beginning with the first elemental position in a random perfect matching, *x*_1_ = 0, we can ask: what is the probability that *x*_1_ is part of a d2-tuple pair (i.e., where each n-tuple has 2 elements)? If the total number of elements *N* is very large relative to the distance threshold *d*, it is highly probable that the position *y*_1_, which matches *x*_1_, is farther than *d* positions away from *x*_1_. If we then choose one of the *d* positions *x*_2_ that are close to *x*_1_ (that is, within the distance threshold *d*), there is a definable chance that its matching position *y*_2_ will also be within the distance threshold *d* of *x*_2_. Specifically, if we circularize the elemental order to avoid boundary effects and ignore overlap of the dn-tuples, for each of the *d* positions following *x*_1_ there are exactly 2*d* possible locations for *y*_2_ (immediately before and after *x*_2_) such that a match would induce the d2-tuple pair {(*x*_1_, *x*_2_), {*y*_1_, *y*_2_}} ∨ *y*_2_ − *y*_1_ ≤ *d*. Note that the use of braces {} and parentheses () around comma-separated lists are used as standard notation to indicate sets that are unordered and ordered, respectively. Therefore, the probability of this happening (still for the moment ignoring boundaries and dn-tuple overlap) is exactly 2 (*d* ∗ *d*)⁄*N* = 2 *d*^2^⁄*N*. If we sequentially update the first position to *x*_1_ = 1,2,3… (*N* − 1), and account for double counting dn-tuples in different orders, the expected number of dn-tuples over the entire dataset is exactly (2 *d*^2^⁄*N*) ∗ *N*⁄2 = *d*^2^. Exact calculations can be made to correct for the added requirements that the paired dn-tuples do not overlap, and that they don’t overlap the end boundary of the dataset, but both of these effects reduce expectations by amounts on the order of multiplying by a factor 1 − *d*^2^⁄*N*; they will both therefore be small if *d*^2^ < *N* and reduce the expected number of dn-tuple pairs compared to the simpler exact circular estimate. The simpler equation is thus conservative, as empirically confirmed by the simulations in Figure 2A. We note that using similar logic, there are about *d*^2^ possible ways for a third matching pair, {*x*_3_, *y*_3_}, to fall within the distance threshold *d* around both members of the d2-tuples for a specific d2-tuple pair, that is {{*x*_1_, *x*_2_, *x*_3_}, {*y*_1_, *y*_2_, *y*_3_}} ∨ *x*_*max*_ − *x*_*min*_ ≤ *d*, *y*_*max*_ − *y*_*min*_ ≤ *d*, where *x*_*max*_ and *y*_*max*_, and *x*_*min*_ and *y*_*min*_, are the respective maximum and minimum integers for each dn-tuple in the pair. Accounting for the dataset size *N*, this happens with a probability close to *d*^2^⁄*N*, albeit with slightly greater reduction factors compared to the earlier case because there are more positions already taken by the members of the specific d2-tuple pair that might be augmented to a d3-tuple pair. Ergo, we expect slightly less than *d*^2^ ∗ (*d*^2^⁄*N*) = *d*^4^⁄*N* d3-tuple pairs in the dataset. Similar reasoning holds for the expected number of d4-tuple pairs, etc., and in general we expect slightly less than *Nd*^2(*n*−1)^⁄*N*^(*n*−1)^ dn-tuple pairs in large datasets with small distance thresholds. This was again empirically confirmed, by the simulations in Figure 2B.

### Updating the null model and calculating false discovery rates

An important step in the analysis of predicted syntenic regions (paired homologous regions of elements with shared possibly collinear genes) is to evaluate the relative likelihood or posterior probability that their order is due to shared descent versus chance arrangement. This can be done in a large variety of ways, but our aim is to estimate, using reasonable assumptions, the probability that paired regions with some number of elements separated by some maximal index distance are truly syntenic; this is the positive predictive values (*PPV*), or conversely the false discovery rate (*FDR*, where *FDR* = 1 − *PPV*). In this manuscript we generally present the *FDR* or *lnFDR* (where *ln* is the natural logarithm) because in many instances the *FDR* is quite small and PPVs near 1.0 are uninteresting to visualize. The simplest and most conservative way to estimate an uncorrected FDR is to assume that all *N* elements are randomly distributed; that is 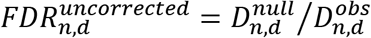, where 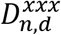is the detected number of paired regions of a particular cluster size *n* within a distance *d*, and *obs* indicates detection in the real dataset, while *null* indicates expected detection under the null model, in this case determined by averages from the Monte Carlo resampling of the real dataset described above. We will often present cumulative FDRs with notation e.g. 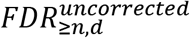, or simply *FDR* if the context is clear from the text and figure or table legends.

The previous analysis, although thorough and relatively simple in mathematical terms, is somewhat dissatisfying for practical interpretation both because the elements that make up observed dn-tuple pairs 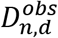 will overlap with the elements that make up e.g. 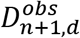, and because the positive identification of matching regions should reduce the proportion of the *N* elements that are randomly assorted under the null model. The first objection is reduced because null expectations are essentially 0 for even moderate cluster sizes under the conditions we studied, and it is not applicable for CMR 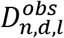 because the lengths *l* of CMRs are the total number of elements in adjacent paired dn-tuples that merged under the specified conditions. If any CMRs overlapped in element constituents in both regions of the pair, they would have been merged, so overlap is not possible.

We chose to address the second objection by serially updating the proportion of the *N* elements thought to be randomly associated under the null model. For CMRs found from a set of dn-tuple pairs, we did this by starting with 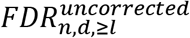 that were less than 0.01 and reduced *N* by the number of elements in both regions of the CMR expected to be truly paired, or 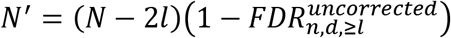. Using this new *N*′, we calculated a new 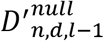 as 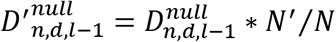, continuing in likewise fashion using the modified *N*′ and 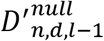 to recalculate the *FDR* for each decremented length down to *l* = *n*. These corrected *FDRs* are designated as 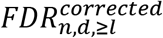, or made clear in the context.

### Kmer matching for rapid protein-coding similarity prediction in seven yeast genomes

To evaluate the effectiveness of this approach to distinguish confident blocks of syntenic, functional elements from disordered, background noise, we evaluated synteny among genomes from seven yeast species in the budding yeast sub-phylum, Saccharomycetaceae. The species used were *Saccharomyces cerevisiae, Saccharomyces paradoxus, Saccharomyces kudriavzevii, Kazachstania africana, Tetrapisispora phaffii, Zygosaccharomyces rouxii* and *Kluyveromyces lactis*. Genomic DNA sequences for all seven species along with their annotations of protein-coding (CDS, or CoDing Sequence) sequences and positions, were obtained from the Hittinger group (Shen et al. 2016; Opulente et al. 2018).

To rapidly create a comparable set of noisy protein homolog predictions among all genomes, a set of reference protein-coding, amino acid sequences from the *S. cerevisiae S288c* genome (obtained from the NCBI database, RefSeq version GCF_000146045.2_R64_protein) was used to scan each genome for matching amino acid kmer elements. This was done by splitting the reference set of protein coding sequences (N=5910) into overlapping fragments of length *k* (e.g., the sequence “MATYPG” kmerized for *k* =3 becomes {“MAT”,“ATY”,“TYP”,“YPG”}). Each set of reference amino acid kmers belonging to each protein coding sequence (CDS, which we often refer to as genes) were stored individually.

Then, each reference CDS kmer set was mapped to the six-frame genome translations (3 codon positions, forward and reverse) of all seven species, including the reference genome, *S. cerevisiae.* Matching regions between the reference kmer sets and six-frame translated kmers in each genome were evaluated in non-overlapping windows, and for a given *k*, a region (a set of adjacent windows) in a query genome was called as containing a potentially homologous protein to a reference CDS based on values for: minimum number of CDS kmer matches for each window in the region, maximum number of windows separating adjacent matching windows, minimum number of adjacent windows containing sufficient CDS matches, and whether or not the genomic kmer matches occurred in the same order as found in the reference CDS (see Table 1). Because these initial regions are in units of windows, and we required no uncertainty in order and match sets, matching regions were resolved to the gene annotation from the Hittinger group (Shen et al. 2016; Opulente et al. 2018) with greatest overlap. We generated three sets of annotations for each of the seven species using three sets of matching criteria, from most conservative or least likely to contain random matches (stringency 1) to the least conservative, stringency 3 (Table 1).

### Mology analysis

The *Mology* algorithm requires ordered integer values as input, so the yeast annotations under a given stringency set were converted from chromosomal coordinates to consecutive integers representing genome order. Annotations for each genome were assigned to a list of ordered integers based on the starting position of the gene annotation from the Hittinger group, regardless of in which reading frame or orientation the match may have been found. In short, the first gene annotated in the first genome was labeled 1, each subsequent gene annotated was labeled with an increment of 1, and the first gene of each consecutive chromosome or genome was labeled one more than the label of the of the last annotated gene in the previous chromosome or genome. After being assigned an order, integer labels from all seven genomes were grouped according to the reference CDS that they matched, such that each reference gene was followed by a list of associated ordered positions across all seven genomes where an annotated CDS match to the reference gene was predicted (See Supplementary Data 1). A map between these ordered identifying numbers and the protein identifiers was retained for back reference after *Mology* analysis (Supplementary Data 1).

*Mology* was run on the consecutive order data of all seven genomes together (dataset Y7G) or on a smaller subset of three genomes (Y3G). Because the groups of matching elements in the yeast dataset were assigned based on reference sequences from *S. cerevisiae*, the original input dataset contained integer positions occurring multiple times, on multiple rows (i.e., protein sequences which matched multiple reference sequences).

As described in the results, *Mology* window size was set to 10 (inline command −d=10) for initial yeast analyses. The window size was set to 2, 5, or 15 in analyses that addressed the effect of window size. The number of matches in dn-tuple pairs was run (inline command −n) as described in the methods for a) −n=2; or b) individually for all values of n from 2..*d*. For the yeast datasets, one hundred pseudoreplicates (inline command −p=100) were generated for each value of *n* to establish a null distribution for dn-tuple pair size and expected frequency. For randomly simulated datasets created for Figure 2, 10,000 replicates were run for *n* = 2 and a wide range of *d* and *N* values, as described in the results.

### Post-Mology analysis of genomic synteny in yeast species

The dn-tuple pairs generated by *Mology* under specified conditions were counted either directly or after merging into CMRs. As described in the Results, the merged regions are simply the merge of all overlapping dn-tuple pairs, and their length was characterized based on the number of matched elements between the two merged regions in a pair. The merging was carried out using a Python script *(cmrs.py*) provided on Github at the same address as above. For some analyses as indicated, we also characterized the total number of annotated genes that were contained within the boundaries of CMRs. For detailed analysis of the regions containing homologues of TOM70/TOM71 in *S. cerevisiae*, we used the reverse map to the reference species (*S. cerevisiae*) to compare to YeastGenomeOrderBrowser syntenic predictions (Byrne and Wolfe 2005) or to the CDS annotations from the Hittinger group (Shen et al. 2016; Opulente et al. 2018).

## Supporting information

Supplementary_Data_Descriptions

## Acknowledgements

We acknowledge the following sources for funding: NIH R01 GM083127 (D.D.P.) and NIH T32 (J.A). We also wish to thank Daniel Polanco for early explorations on this topic.

