## Supplementary_Figures for "A fast, general synteny detection engine"

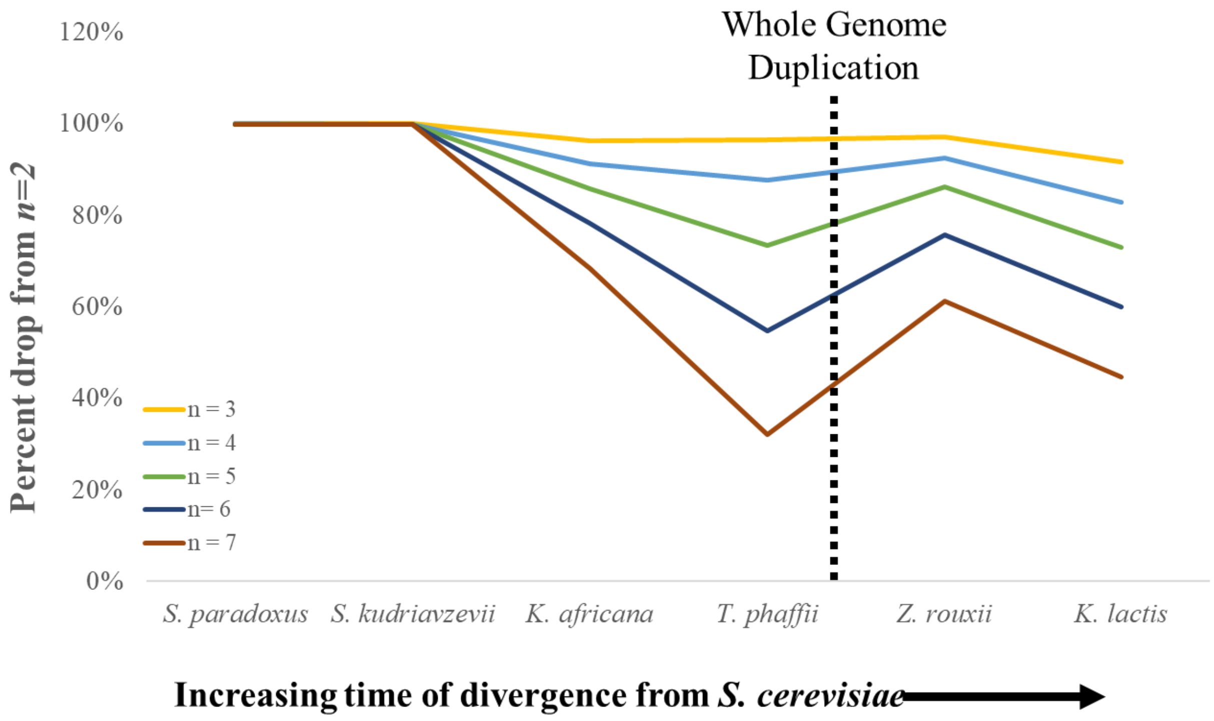


**Supplementary Figure 1. Decay of genes in predicted syntenic regions with *S. cerevisiae* depending on dn-tuple size.** The number of genes shown are contained in CMRs between *S. cerevisiae* and six other species from across the budding yeast sub-phylum, Saccharomycetaceae. The percent drop off compared to Mology analyses with n=2 is shown for n=3 to n=7. The data and axes labels for this figure are the same as in Figure 5.

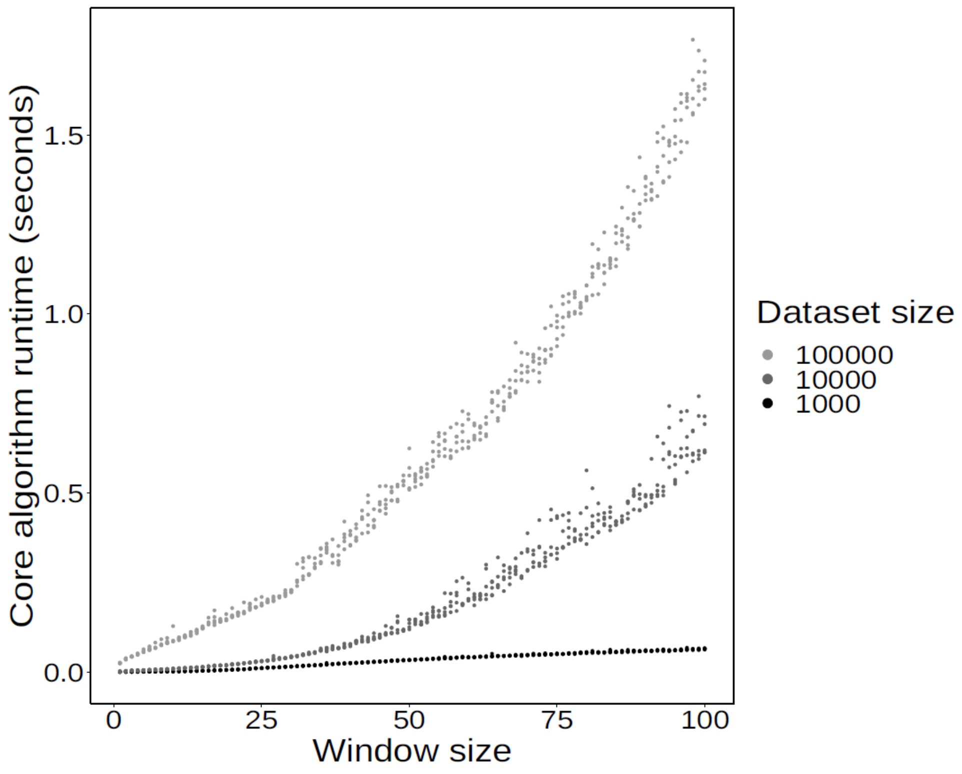


**Supplementary Figure 2. Predicted versus simulated dn-tuple pairs.** Core *Mology* algorithm run time for simulations in Figure 2A of main text. Shading of points indicates the number of randomly-paired elements in each simulated dataset (100 K, 10 K, or 1 K), as indicated in legend. Window size (*d*) is positively correlated with run time, and all conditions were analyzed in under 2.0 seconds.
