## Supplementary_Data_Descriptions for "A fast, general synteny detection engine"

### Supplementary Data and Supplementary Tables

*In order of appearance*

**Supplementary Data 1. The sets of all genes with high similarity to S. cerevisiae protein-coding sequences, at three stringency levels**. Yeast kmer annotations. This is a readme plus a folder for each stringency, along with a folder labeled "yeast genomes", which contains the yeast genome sequences and protein-coding .gtf files from the Y1000+ project generated by the Hittinger group (Opulente, *et al.*, 2018). Files are *_merged.txt, which contains *S. cerevisiae* (Scer) ref IDs that mapped to a given genome, ranked by order of appearance in that genome. The ones on the same lines are multiple Scer proteins that mapped to a single Y1000+ gene annotation. The "*_stringent1.txt" files are raw output that contain Scer ref protein IDs (and full name information), followed by square bracket sets of three numbers in parentheses, e.g. (23.0, 43173, 67) which represent: (relative position in match window, genomic nucleotide coordinate of match, position of match in ref CDS); at the end of the square bracket is the translation frame.

**Supplementary Data 2. Supplementary Data 1, organized into the groups that match each S. cerevisiae protein-coding sequence.** Mology input. These are the order maps at three stringencies. Each stringency has an associated _merged.txt, _merged_speciesBounds.txt, and merged_noIDs.txt. No_IDs are just the NP numbers converted to their rank number in the overall dataset; this is the input file for *Mology*. The species bounds files provide the first and last rank number for each genome.

**Supplementary Data 3. Mology-detected dn-tuples at $d = 10$ for values of $n$ from 2-10.** Contains *_raw.txt and *_evalue_table.txt, currently all compressed together, for Y7G, all three stringencies. Raw is the ordered integer sets and lengths of dn-tuple pairs, whereas evalue table are the counts of real genomic dn-tuple pairs and means across 100 pseudoreplicates. Raw for example contains

- {[0 3 4 5 6] 6 [76 84] 8 8}
- original {[1 4 5 6 9] 8 [80 89] 9 9}

or

- {[9102 9103] 1 [28391 28392] 1 1}
- original {[9275 9276] 1 [28564 28565] 1 1}

This is direct Mology output. The first line in the pair represents the rank-ordered integers implemented by *Mology*. The second line represents the original position of the matching genes in their respective genome. The indices in the 'original' record may be greater than in the rank-ordered *Mology* record due to the exclusion of singletons in the S. cer reference genomes. The number of matching elements in each half of the pair is the number of integers in each set of brackets. The integers in each bracket pair represent the matching genes in each genome. The integers in between brackets represent the distance between beginning and end of each dn-tuple and the last number is the longer dn-tuple length of the dn-tuple pair.

The evalue tables contain e.g. for n6 d10:

- group_size real_count mean_p_rep_count

- 1 0 0
- 2 0 0
- 3 0 0
- 4 0 0
- 5 0 0
- 6 40259 0
- 7 0 0

For dn-tuples generated under the “strict” setting, *Mology* only outputs the dntuple pair counts for a given value of n at a time.

**Supplementary Data 4. Mology-detected dn-tuples at $d = 10$ for values of $n$ from 2-10, pseudoreplicates.** This has each stringency tarred separately, containing a separate text file for each pseudoreplicates, from 0-99.When files are empty, no dn-tuples were generated for that pseudoreplicate.. Non-empty files have paired lines, such as:

- {[1869 1872 1876 1878] 9 [24872 24874 24881 24882] 10 10}
- original {[1927 1930 1934 1936] 9 [25045 25047 25054 25055] 10 10}

This is direct Mology output. The first line in the pair represents the rank-ordered integers implemented by *Mology*. The second line represents the original order of the matching genes in their respective genome. The number is somewhat reduced, due to the exclusion of singletons in the Scer reference genomes. The number of matching elements in each half of the pair is the number of integers in each set of brackets. The integers in each bracket pair represent the matching genes in each genome. The integers in between brackets represent the distance between beginning and end of each dn-tuple and the last number is the longer dn-tuple length of the dn-tuple pair.

**Supplementary Table 1. Annotation match counts for *S. cerevisiae* reference protein-coding sequences in Y7G and Y3G at three stringencies**. This file contains two tabs: one for the S. cer reference annotation counts of Y7G and one for the Scer reference annotation counts of Y3G, at all three stringencies.

**Supplementary Table 2. CMR length counts for Y7G and pseudoreplicates at three stringencies**. Contains the distributions of CMR lengths (number of genes per a given CMR) and the frequency with which a CMR of that size occurs in each genome in Scer vs. species comparisons. The CMRs represented in these tables were generated from the following settings: Stringency 1, n=3, d=10. Stringency 2, n=4, d=10. Stringency 3, n=6, d=10.

**Supplementary Data 5. Mology-detected CMRs at $d = 10$ for values of $n$ from 2-10.** These are for Y7G, and there is a file for each n and stringency, although stringency 1 is called “strict”, and stringency 3 doesn’t come in until n=5. An example line is:

- [[10, 15, 18, 19], [5381, 5382, 5387, 5391]]

These are the results of cmrs.py, where dn-tuples have been merged into CMRs. These contain the original genome integers of matching genes. These can be used to reverse map back to the proteinID.

**Supplementary Data 6. Mology-detected CMRs at $d = 10$ for values of $n$ from 2-10, Y7G pseudoreplicates.** CMR Stringency 1, n=2 to 5, stringency 3 n=6-8.

- [[10, 15, 18, 19], [5381, 5382, 5387, 5391]]

These are the results of cmrs.py, where dn-tuples have been merged into CMRs. These contain the original genome integers of matching genes. These can be used to reverse map back to the proteinID

**Supplementary Table 3. FDR correction**. The FDR calculations that are the source of Figure 3. Calculated for stringency 1, d=10, n=2. Compares *S. cerevisiae* CMR lengths to the null.

**Supplementary Table 4. Dn-tuple and CMR pair counts for Y3G and pseudoreplicates at three stringencies, n=2-15, d=5, 10 and 15.** There is a different sheet for each d value. These are summaries of the data in Supplementary Data 7 (below)..

**Supplementary Data 7. Mology-detected dn-tuples and CMRs, for Y3G and Y3G pseudoreplicates, d=5,10,15, n=2-15.** Contains all the Mology dn-tuple and CMR pair output for each value of d at each value of n, from the Y3G dataset. This includes the corresponding data for the Y3G pseudoreplicates, 0-99. At all three stringencies. Data is grouped into separate folders by d= and stringency=.

**Supplementary Table 5. TOM region analysis results.** There are 5 species comparisons to Scer, each of which contains 3 different stringencies, d=10 and n=3, 4, or 6.Shown are the CMRs that contain the example S. cer paralogs of interest (TOM70/TOM71), between Scer and each species. Each CMR contains the matching S. cer protein identifiers (NP_ID) found in that region between Scer and each species..

**Supplementary Table 6. Number of protein IDs assigned to same genomic position as given NP_ID in the TOM regions.** This shows a single table, comparing each stringency for Zrou and Klac. The frequencies at each stringency represent how many putative homologs matched to each S. cer NP_ID.

**Supplementary Table 7. Gene counts in CMRs for all pairwise species comparisons, n=2 to n=10.** This shows the gene counts for each comparison at stringency 1, for CMR pairs between all pairwise species combinations. These are counts of genes contained within CMRs in a given genome, relative to its comparison species. d=10.

**Supplementary Table 8. Gene counts in CMRs for only *S. cerevisiae* pairwise species comparisons., n=2 to n=10.** The counts of genes in CMRs, across three stringencies, and across all 6 comparative species. Values of n start at differing settings for each stringency. This is the data that Figure 5 is generated from. And the individual counts are included in SuppTable 7

All information, including supplementary tables and supplementary figure legends, must be uploaded separately as "supplementary files" and not placed in the manuscript.

Files submitted as supplementary material must be correct and complete, including figure legends, etc. These files will not be edited or reformatted, but are posted exactly as submitted.

The following formats are acceptable: Plain text (.txt); HTML (.html, htm); Jpeg (.jpg, .jpeg); GIF (.gif); QuickTime video (.mov); MPEG Movie (.mpg); Microsoft AVI Video (.avi); Adobe PDF (.pdf); Microsoft Excel Spreadsheet (.xls)
